# Metabolic remodelling in hiPSC-derived myofibres carrying the m.3243A>G mutation

**DOI:** 10.1101/2024.06.11.598565

**Authors:** Gabriel E. Valdebenito, Anitta R. Chacko, Chih-Yao Chung, Preethi Sheshadri, Haoyu Chi, Benjamin O’Callaghan, Henry Houlden, Hannah Rouse, Valle Morales, Katiuscia Bianchi, Francesco Saverio Tedesco, Robert D. S. Pitceathly, Michael R. Duchen

## Abstract

Mutations in mitochondrial DNA cause severe multisystem disease, frequently associated with muscle weakness. The m.3243A>G mutation is the major cause of Mitochondrial Encephalomyopathy Lactic Acidosis and Stroke Like episodes (MELAS). Experimental models that recapitulate the disease phenotype *in vitro* for disease modelling or drug screening are very limited. We have therefore generated hiPSC-derived muscle fibres with variable heteroplasmic mtDNA mutation load without significantly affecting muscle differentiation potential. The cells are excitable and show physiological characteristics of muscle fibres and show well organised myofibrillar structure. In cells carrying the m.3243A>G, the mitochondrial membrane potential and oxygen consumption were reduced in relation to the mutant load. We have shown through proteomic, phosphoproteomic, and metabolomic analyses that the m.3243A>G mutation variably affects the cell phenotype in relation to the mutant load. This variation is reflected by an increase in the NADH/NAD^+^ ratio, which in turns influences key nutrient-sensing pathways in the myofibres. This model enables detailed study of the impact of the mutation on cellular bioenergetics and on muscle physiology with the potential to provide a platform for drug screening.

## Main

Mitochondrial myopathies are mitochondrial diseases usually caused by mutations of nuclear or mitochondrial encoded proteins and characterised by muscle weakness. The m.3243A>G DNA mutation typically causes a disease known as MELAS – Mitochondrial Encephalomyopathy Lactic Acidosis and Stroke Like episodes. Muscle weakness is a major feature and can be profoundly disabling. The biochemical and physiological consequences of this specific mutation in terminally differentiated tissues remain poorly understood. Cells carrying the m.3243A>G mutation exhibit heteroplasmy, defined as the presence of both wild-type and mutant mitochondrial DNA (mtDNA). Broadly speaking, disease severity maps to mutant load, although the relationship between genotype and phenotype is poorly understood. Currently no effective disease modifying treatments are available. A major hurdle in understanding the pathophysiology of the disease and in finding treatments is the lack of good experimental systems for disease modelling or for drug screening. Currently, there are limited tools available for generating models of pathogenic mtDNA mutations. Gene editing has not yet evolved to the point we can engineer the mitochondrial genome to produce animal models with specific mitochondrial DNA (mtDNA) mutations (Silva-Pinheiro & Minczuk, 2022). We have therefore generated human induced pluripotent stem (hiPS) cells from patient derived fibroblasts carrying the m.3243A>G mutation and in the present paper, we describe the generation of viable muscle fibres from the hiPS cells by recapitulating key signaling events during myogenesis (Al Tanoury et al., 2021; Chal & Pourquié, 2017).

### Implementing an *in vitro* model to study the m.3243A>G mutation

One of the key features of many mtDNA diseases is the expression of heteroplasmy. We took advantage of the random segregation of mtDNA and manually selected clones, which were then expanded and maintained as stable hiPSC lines expressing variable burdens of mutant mtDNA (Fig. 1a, b). This has the added advantage of enabling the generation of isogenic stem cell lines - a significant advantage in investigating the disease phenotype. Three different cell lines were established, bearing 50% (hiPSC-M50), 90% (hiPSC-M90), and undetectable levels (hiPSC-M0) of the m.3243A>G mutation (Fig. 1c). These cell lines exhibited no differences in terms of pluripotency, as all colonies were positive for nuclear markers such as SOX2, NANOG, OCT4, and the surface marker SSEA4 (Supplementary Fig. 1a, b). Measurements of gene expression levels through qPCR confirmed the expression of these markers, in contrast to an unrelated human fibroblast line (Supplementary Fig. 1c). The colonies did not exhibit significant differences in terms of oxygen consumption (Fig. 1d), measured using the ‘Seahorse’ assay, or mitochondrial membrane potential (Δψm), measured using the equilibration of Tetramethylrhodamine, methyl ester (TMRM, Fig. 1e, Fig. 1f and Supplementary Fig. 1d).

**Figure 1.**
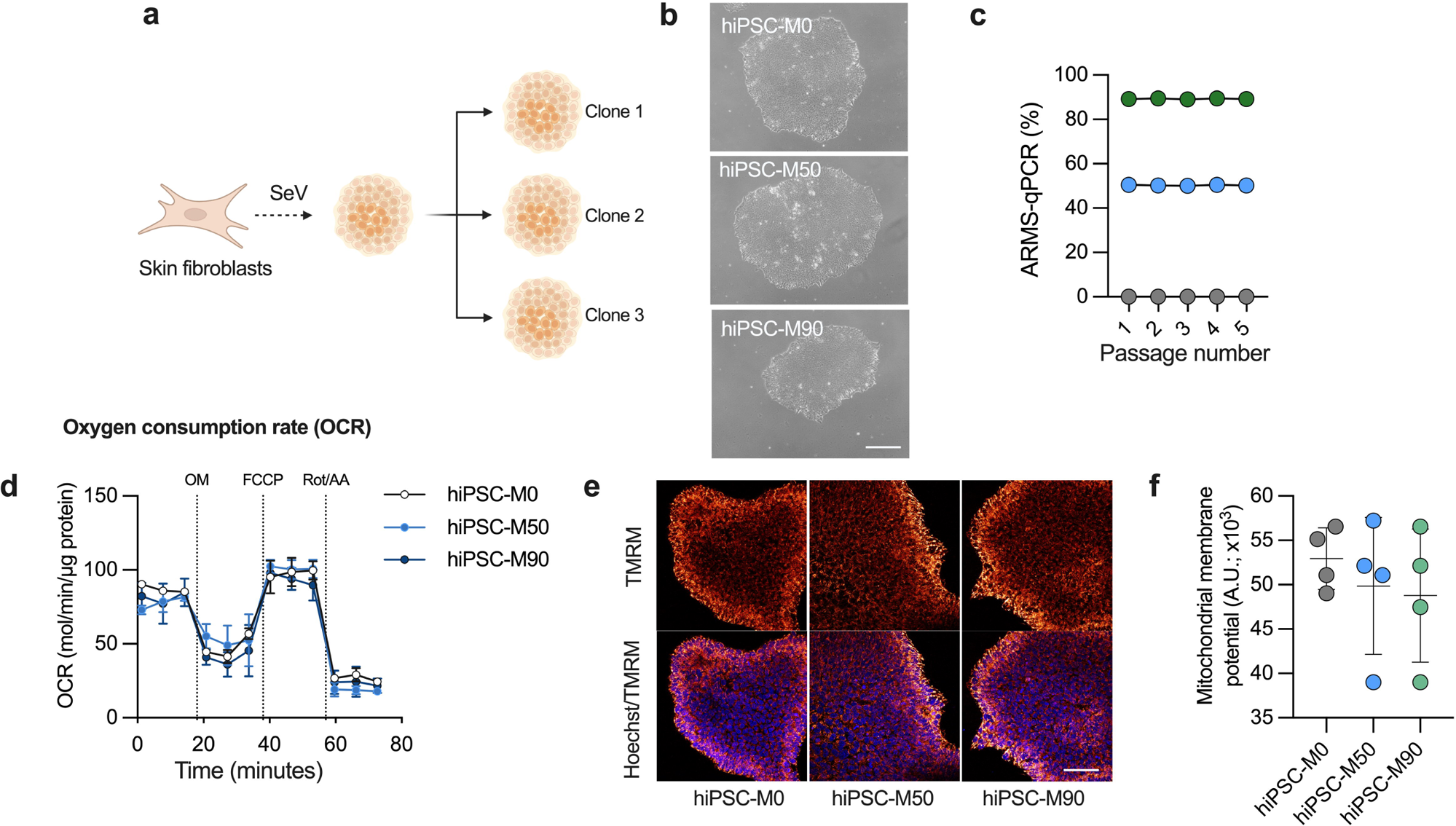
The m.3243A>G mutation remains stable in hiPSCs and does not affect pluripotency or mitochondrial function. (A) Schematic showing experimental approach to select hiPSC colonies carrying different levels of the m.3243A>G mutation. Created with BioRender.com (B) Bright-field micrographs of hiPSC colonies in feeder-free conditions. Scale bar = 100 µm. (C) Mutant load quantification through ARMS-qPCR in hiPSC clones. *n* = 3 independent biological samples). (D) Cell respiratory capacity measured using the Seahorse XFe96 extracellular flux analyser in hiPSC colonies. *n* = 3 independent biological samples, 5 culture wells per cell line. (E) Confocal images of hiPSC loaded with 25 nM Tetramethylrhodamine, methyl ester (TMRM) and 1µg/mL Hoechst 33342. Scale bar = 150 µM. (F) Averaged quantification mutant load of hiPSC colonies loaded with TMRM (n = 3 independent biological samples). Source data are provided as a Source Data file. All data were represented as mean ± SD and data were analysed by one-way ANOVA with Tukey’s multiple comparisons test (*p < 0.05, **p < 0.01, ***p < 0.001, ****p < 0.0001).

### hiPSC bearing the m.3243A>G can successfully differentiate into mature myofibres

To establish a muscle model that accurately reproduces key characteristics of muscle myopathy in carriers of the m.3243A>G mutation, we employed a differentiation protocol designed to recapitulate crucial signaling events in muscle development. The differentiation strategy involves a combination of FGF2 and the Wnt agonist CHIR99021 to drive cell commitment towards the mesodermal lineage, while simultaneously inhibiting bone morphogenetic protein (BMP) signaling with LDN193189 to restrict mesodermal fates to the presomitic mesoderm (Diaz-Cuadros et al., 2023; Diaz-Cuadros et al., 2020) (Fig. 2a). The induction of all three cell lines was initially assessed at day 7 of differentiation, during which clusters of cells (myocenters) were present in the preparation, surrounded by muscle progenitors (Supplementary Fig. 1e). From day 12 onward, the media were modified by replacing CHIR99021 and LDN193189 with growth factors (HGF and IGF) to facilitate the expansion of myogenic progenitors. These progenitors were subsequently replated to establish a highly homogeneous cell culture, which was then induced to differentiate into mature myofibres in a process called secondary myogenesis.

**Figure 2.**
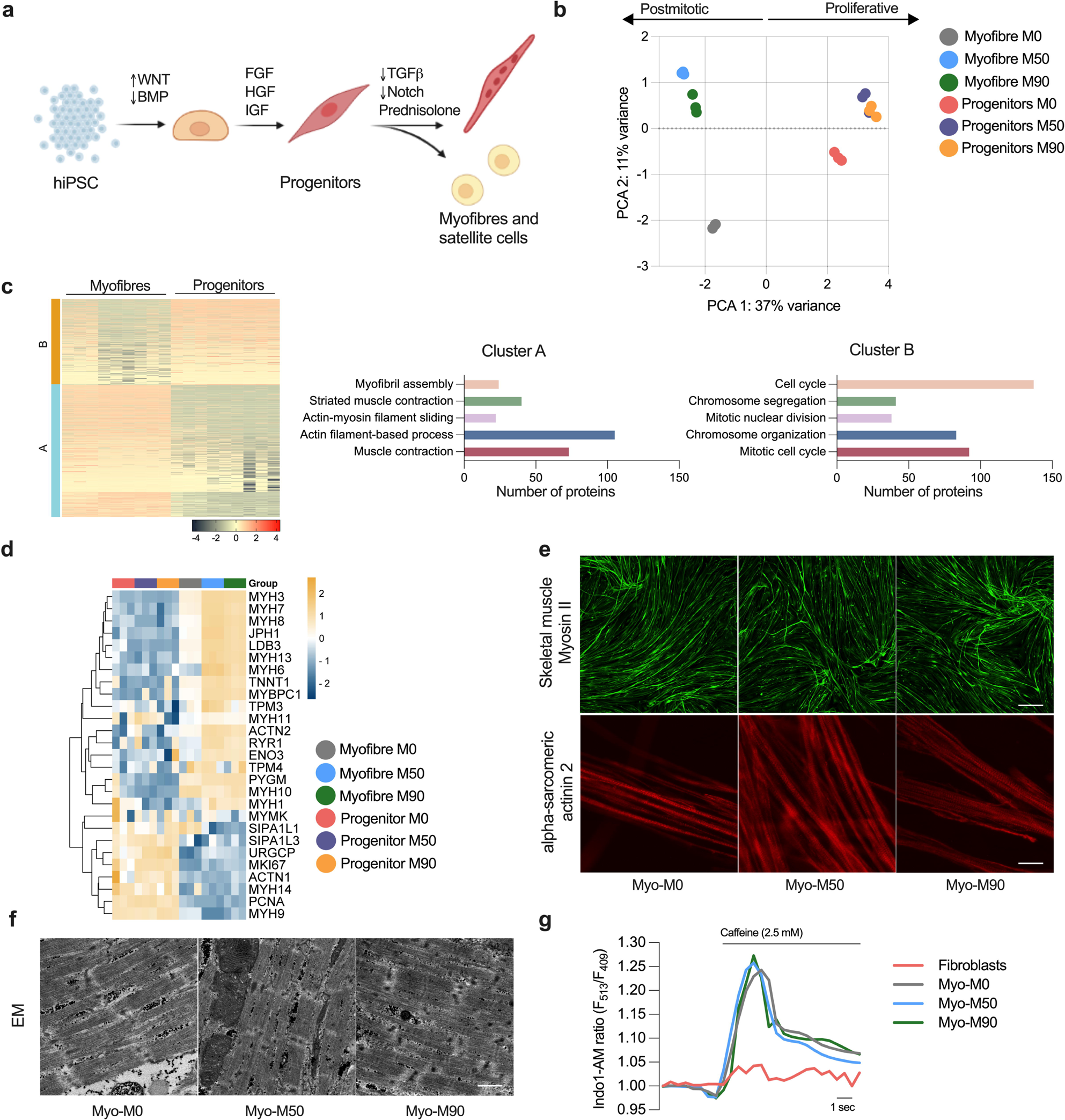
Generation of myofibres derived from hiPSC bearing the m.3243A>G. (A) Protocol used to direct the cells into the mesodermal fate and terminal differentiation of muscle progenitors. (B) Principal component analysis (PCA) of protein signature showing variance between sample groups (n = 3 replicates per condition). (C) K-means clustering heat map of proteins (left, n = 3 replicates) and quantification of the top hits in both clusters. (D) Heatmap representing proteins associated with muscle maturation in myofibres and protegnitors (n = 3 replicates). (E) Representative confocal images of myofibres stained with antibodies against Skeletal Muscle Myosin (top, scale bar = 500 µM) and α-Sarcomeric Actinin 2 (bottom, scale bar = 20 µm). (F) Electron micrograph of myofibres sections. Scale bar = 1 µM. (G) Representative changes in Indo-1 AM fluorescence intensity after stimulation with 2.5 mM Caffeine.

To closely examine this transition and identify potential deviations at the progenitor level caused by the presence of the m.3243A>G mutation, which could potentially interfere with the final differentiation process, we conducted a comparative proteomic analysis between the progenitor cells and myofibres, both with and without the mutation. The score plot of principal component analysis (PCA) revealed distinct clustering of progenitors and myofibres in separate quadrants of the plot (along PC1), indicating notable differences in these two cell types. Progenitors represent a proliferative cell type, whereas muscle cells are postmitotic and possess a contractile apparatus that is absent in the progenitor population. In the context of the mutant lines, it is noteworthy that the progenitors continued to exhibit close clustering. However, there was a noticeable increase in variance observed between the isogenic control and mutant myofibres (separated along PC2). This heightened variance suggests that the differences in the disease phenotype are significantly more pronounced in postmitotic cells compared to the progenitors (Fig. 2b), a cell type less dependent on oxidative phosphorylation (Sin et al., 2016).

We also employed K-means clustering algorithm to gain insights into the underlying structure of the dataset (Fig 2c). To elucidate the biological significance of these clusters, we conducted Gene Ontology (GO) analysis. As expected, the top cluster, enriched in progenitors (cluster B), contained GO terms associated with proliferative cells, such as the cell cycle, chromosome segregation, and organization, as well as mitotic nuclear division. Meanwhile, the bottom cluster (cluster A), highly enriched in myofibres, exhibited a significant enrichment in GO terms related to myofibril assembly, striated muscle contraction, actin filament sliding, and associated processes.

Based on previously reported protein expression profiles in mature myofibres and progenitors, we curated a list of muscle-related proteins to observe their expression across samples. As expected, isoforms of myosin heavy chain were highly abundant in all myofibres compared to the progenitors. Although not all these myosin isoforms are skeletal muscle-specific, as some are also expressed in smooth muscle. However, MYH3, 7, and 8 (the top hits) belong to proteins associated strictly with skeletal muscle specification and maturity (Fig 2d). Junctophilin-1 (JPH1), which contributes to the construction of the skeletal muscle triad by linking the t-tubule to the sarcoplasmic reticulum, was also highly enriched in myofibres when compared to progenitor cells. LDB3, TNNT1, RYR1 and ACTN2 were also enriched in the differentiated cells, confirming that all components of the muscle machinery expected in a differentiated myofibre were expressed in the preparation. It is important to note that a few of these proteins were more abundant in the mutant lines compared to the isogenic control. However, when considering all three lines together, their expression levels were higher than in the progenitors.

We also incorporated proteins associated with cell proliferation into this curated list. Since muscle cells are postmitotic, the expression of these proteins is expected to be lower compared to the progenitors, which are proliferating. The signal-induced proliferation-associated 1/3-like proteins (SIPA1L1 and SIPAL1L3), the upregulator of cell proliferation (URGCP), and the proliferating cell nuclear antigen (PCNA) were all upregulated in the progenitors. One of the common markers of proliferation, MKI67, was also highly enriched in the progenitors and downregulated across all the myofibres.

To understand the impact of the m.3243A>G mutation on muscle structure, we stained fixed preparations of myofibres with an antibody against the heavy chain of myosin II, specifically targeting the light meromyosin portion. Through confocal imaging, we observed positive staining of the myofibres in all preparations (Fig 2e, top panel). Furthermore, we labelled the myofibres with an antibody against alpha-actinin 2, a structural protein expressed in both skeletal and cardiac muscles that serves to anchor myofibrillar actin thin filaments and titin to Z-discs. All fibres were positive for alpha-actinin 2, and showed striations (Fig 2e, bottom panel). To observe the ultrastructure of myofibres more closely, cells were examined by electron microscopy, revealing the presence of sarcomeres along the muscle fibres in all three preparations with no appreciable differences between the different cell lines (Fig 2f). Alpha-actinin staining was pseudo-coloured based on local fiber orientation. Assessing the spatial organisation of neighbouring myofibers revealed aligned bundles, despite the absence of predefined orientation cues (Supplementary Fig. 2a). This observation implies the existence of a self-organizing process during myofiber bundling, a characteristic trait of mature preparations (Mao et al., 2022).

Muscle contraction is driven by the cytosolic calcium signal which in turn is shaped by mitochondrial function - both in terms of energy supply and mitochondrial calcium uptake. To test whether the muscle cells were excitable, we first loaded the preparations with the ratiometric calcium indicator Indo-1 AM. All three cell lines showed a response to stimulation with 2.5 mM Caffeine, with an increase in cytosolic calcium concentration. There was no difference in the response to caffeine between the isogenic control and mutant myofibres. This response is dependent on the expression of RyR3 Ca^2+^ release channels in the sarcoplasmic reticulum (SR) and no response was measurable in fibroblasts (Fig 2g). Electrical excitability was confirmed in all differentiated myofiber lines, which showed brisk calcium transients in response to electrical stimulation with no appreciable differences (Supplementary Fig. 2b). These findings show that cell reprogramming is not significantly impaired by the m.3243A>G mutation and the mutant cells exhibit physiological response to both chemical and electrical stimulation expected from differentiated muscle fibres.

### Mitochondrial bioenergetic is impaired in myofibres carrying the m.3243A>G mutation

Mitochondrial dysfunction is commonly considered a potential hindrance to cellular differentiation as it often impacts development (Qi et al., 2022). To evaluate whether myogenic specification was altered in the *in vitro* model, we quantified the myogenic efficiency by calculating the percentage of nuclei within α-actinin 2 positive cells compared to the total number of nuclei (Fig. 3a). No appreciable differences were observed between the isogenic control and mutant myofibres. We also measured the length and width of myofibres within the preparations. While there were no significant differences in these variables among the three lines during terminal differentiation, mean cell diameter at day 10 was decreased in myo-M90 cells. (Fig. 3b). The ratio of total protein concentration to genomic DNA was decreased in both mutant lines compared to control (Fig. 3c). Global protein synthesis, measured by the incorporation of puromycilated peptides (Fig. 3d), was also decreased in the mutant lines. The proteomic analysis also showed enriched terms related to muscle-associated conditions, with ‘abnormality of the musculature’, ‘abnormal muscle physiology’, and ‘skeletal muscle atrophy’ as the top hits (Fig. 3e).

**Figure 3.**
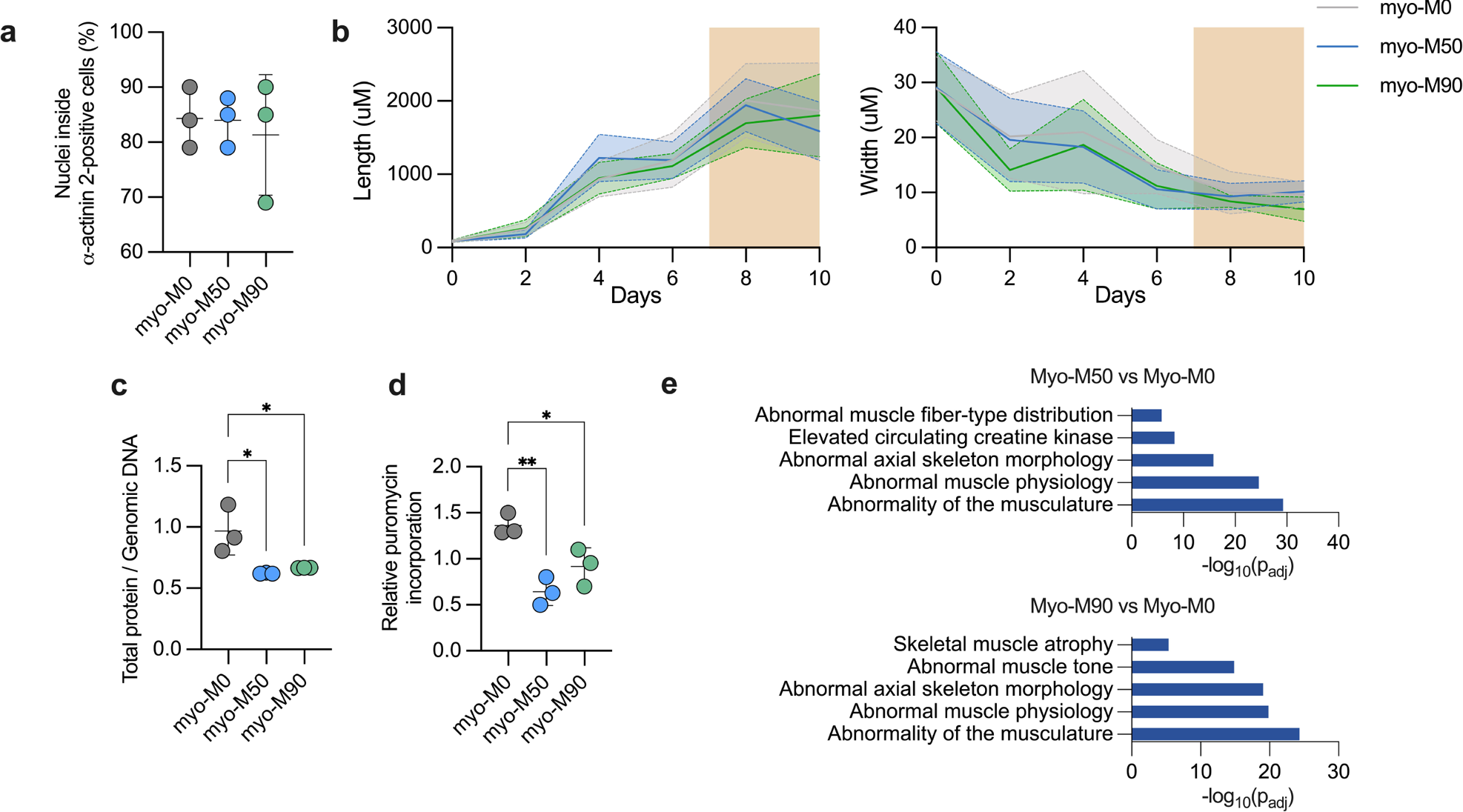
Myogenic efficiency and muscle phenotype in cells carrying the m.3243A>G mutation. (A) Quantification of myogenic differentiation. Nuclei outside and inside α-actinin 2-positive cells are counted and then the ratio of nuclei inside α-actinin 2-positive cells / total nuclei are used to calculate the differentiation efficiency. Results are expressed as percentage of the total population in the culture (n = 3 independent biological replicates). (B) Length and width of myocytes, myotubes and myofibres over a period of 10 days of differentiation. Dotted and solid lines show the mean length and width of each day, while shaded area shows standard deviation (n = 3 independent biological replicates). (C) Total cellular protein content relative to genomic DNA in myotubes (n = 3 independent biological replicates). (D) Relative translation rate expressed as puromycin incorporation (n = 3 independent biological replicates). (E) Human Phenotype Ontology of the differentially expressed proteins between Myo-M50 vs Myo Myo-M0 (top) and Myo-M90 vs Myo-M0 (bottom). Source data are provided as a Source Data file. All data were represented as mean ± SD and data were analysed by one-way ANOVA with Tukey’s multiple comparisons test (*p < 0.05, **p < 0.01, ***p < 0.001, ****p < 0.0001).

To explore the metabolic impact of the m.3243A>G mutation, we first carried out ARMS qPCR to ensure the mutant load was retained after differentiation. ARMS qPCR measurements established that mutant load did not vary over time in the different cell lines (Fig. 4a). Respiratory rate was measured using the Seahorse XFe96 extracellular flux analyser. Both basal and maximal uncoupler-induced oxygen consumption rates were significantly reduced in both mutant lines (Fig. 4b). To establish that these observations in mitochondrial respiration are common for the m.3243A>G mutation, we differentiated an unrelated hiPS cell line carrying a 70% m.3243A>G mutant load detected by ARMS-qPCR (Supplementary Fig. 4a) following the same protocol (Fig. 2a) and compared the data against a control hiPSC line derived from a healthy donor. Basal and maximal respiratory rates were also reduced in this mutant cell line (Supplementary Fig. 4b).

**Figure 4.**
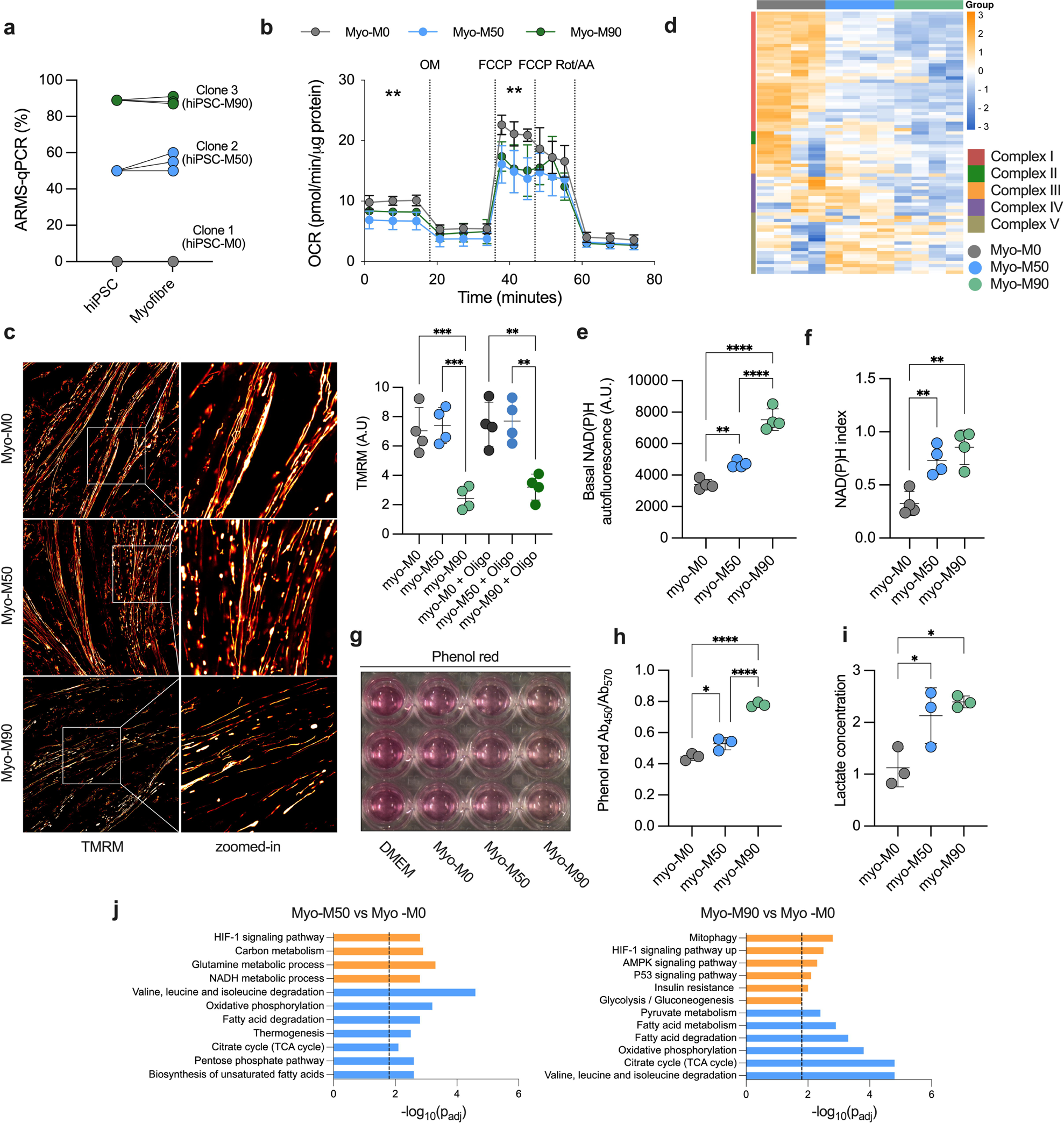
Myofibres expressing the m.3243A>G show mitochondrial dysfunction. (A) Changes in mutation load from hiPSC to fully differentiated myofibres (n = 3 independent biological replicates). (B) Cell respiratory capacity measured using the Seahorse XFe96 extracellular flux analyser in myofibres normalised by protein concentration (n = 3, 6 culture wells per experiment). (C) Confocal images of myofibres loaded with 25 nM Tetramethylrhodamine, methyl ester (TMRM, left) and quantification of mitochondrial membrane potential (right). n = 4 independent biological replicates. (F) Quantification of lactate concentration in the media, normalised by protein concentration. n = 5 independent biological replicates. (E) Levels of basal mitochondrial NAD(P)H measured by NAD(P)H autofluorescence, A.U: arbitrary units. n = 3 independent biological replicates. (F), Quantification of NADPH redox index. n = 4 independent biological replicates. (G) Representative image of culture wells showing a change in the media colour. (H) Absorbance ratio of phenol-red. n = 3 independent biological replicates. (I) Fold-changes in lactate concentration measured by CuBiAn. n = 3 independent biological replicates. (J) Top upregulated (orange) and downregulated (blue) KEGG pathways from the proteomic dataset. Source data are provided as a Source Data file. All data were represented as mean ± SD and data were analysed by one-way ANOVA with Tukey’s multiple comparisons test (*p < 0.05, **p < 0.01, ***p < 0.001, ****p < 0.0001).

Meanwhile, the Δψm was significantly decreased only in the line carrying the highest mutant load (M90; Fig. 4c). To ask whether the Δψm in Myo-M50 is conserved by reversal of the F_1_F_o_-ATP synthase we treated the cells with oligomycin. In cells in which the ATP synthase works ‘in reverse’ (i.e. as a proton translocating ATPase), oligomycin causes a collapse of the membrane potential. In the Myo-M50 cells, oligomycin did not alter the Δψm, showing that the potential is not maintained by the reversal of the ATP synthase (McKenzie & Duchen, 2016; Valdebenito, Chacko, & Duchen, 2023). Expression levels of subunits of complex I and II (NDUFB and SDHB) of the electron transport chain (ETC) were decreased in response to the m.3243A>G mutation when measuring representative proteins of the ETC (Supplementary Fig. 4c). To observe global changes in mitochondrial protein expression in Myo-M50 and Myo-M90, we compared the mitochondrial proteome of the ETC components. Notably, we observed that expression of most of the proteins in complex I were significantly reduced in both mutant lines. This observation was even more pronounced in Myo-M90, where complexes II, III, and IV were also downregulated. Even though complex II is entirely encoded by the nuclear genome we have noted before in fibroblasts carrying the m.3243A>G mutation that complex II expression was reduced (Chung et al., 2021). Moreover, complexes IV and V were upregulated in Myo-M50 when compared to both the isogenic control and Myo-M90 (Fig 4d).

Complex I is the component of the ETC which oxidises NADH. Given a decrease in the expression of complex I subunits, we took advantage of the intrinsic fluorescence of NADH to measure the ability of complex I to oxidise this molecule. Under confocal UV excitation, we quantified the basal mitochondrial NADH autofluorescence (Fig 4e and Supplementary Fig. 4d). As expected, both mutant lines showed increased NADH autofluorescence, with a more prominent increase in Myo-M90, consistent with the elevated mutant load and more severely impaired function (Fig. 4e). In order to measure changes in the redox state, we treated the cells with NaCN and FCCP to obtain the maximal (fully reduced) and minimal (fully oxidised) autofluorescence signals respectively. The autofluorescence signal normalised between these values confirmed a more reduced state of the NADH/NAD+ pool in both mutant lines compared to the isogenic control (Fig. 4f).

We observed that the colour of phenol red, a pH indicator in the growth media, was more yellow in both cultures of mutant lines compared to the control (Fig. 4g, h), suggesting acidification consistent with an elevated lactate concentration. This was confirmed when measured using the CuBiAn HT-270 (Fig. 4i), recapitulating the lactic acidosis seen in patients with MELAS. GO analysis also suggested the downregulation of tricarboxylic acid (TCA) cycle and Oxidative Phosphorylation in Myo-M50 and increased glycolysis in Myo-M90, pointing to the reprogramming of these metabolic pathways as a consequence of the m.3243A>G mutation (Fig. 4j). Together, these data show that the m.3243A>G mutation alters the bioenergetics of the myofibres, affecting the expression of ETC subunits and promoting increased glycolysis.

### Rewiring of nutrient signaling pathways is accompanied by increase NADH/NAD **^+^** ratio and compensatory changes in NADH shuttles

In the cytosol, the increased rate of glycolysis and NADH generation leads to the conversion of pyruvate into lactate via lactate dehydrogenase, which restores NAD+ levels and increases lactate concentration. We quantified the lactate-to-pyruvate ratio reflecting the cytosolic NADH/NAD^+^ ratio, which was increased in the mutant lines, with a consistent upregulation in the line carrying the higher mutant load (Fig. 5a). We corroborated this observation with the genetically encoded cytosolic NADH/NAD+ sensor, Peredox, which further confirmed these findings (Fig. 5b). Targeted metabolomic analysis also showed changes in the labelled pattern of some metabolites, especially α-Ketoglutarate, which is involved in the conversion of NAD^+^ to NADH in the TCA cycle (Supplementary Fig. 5).

**Figure 5.**
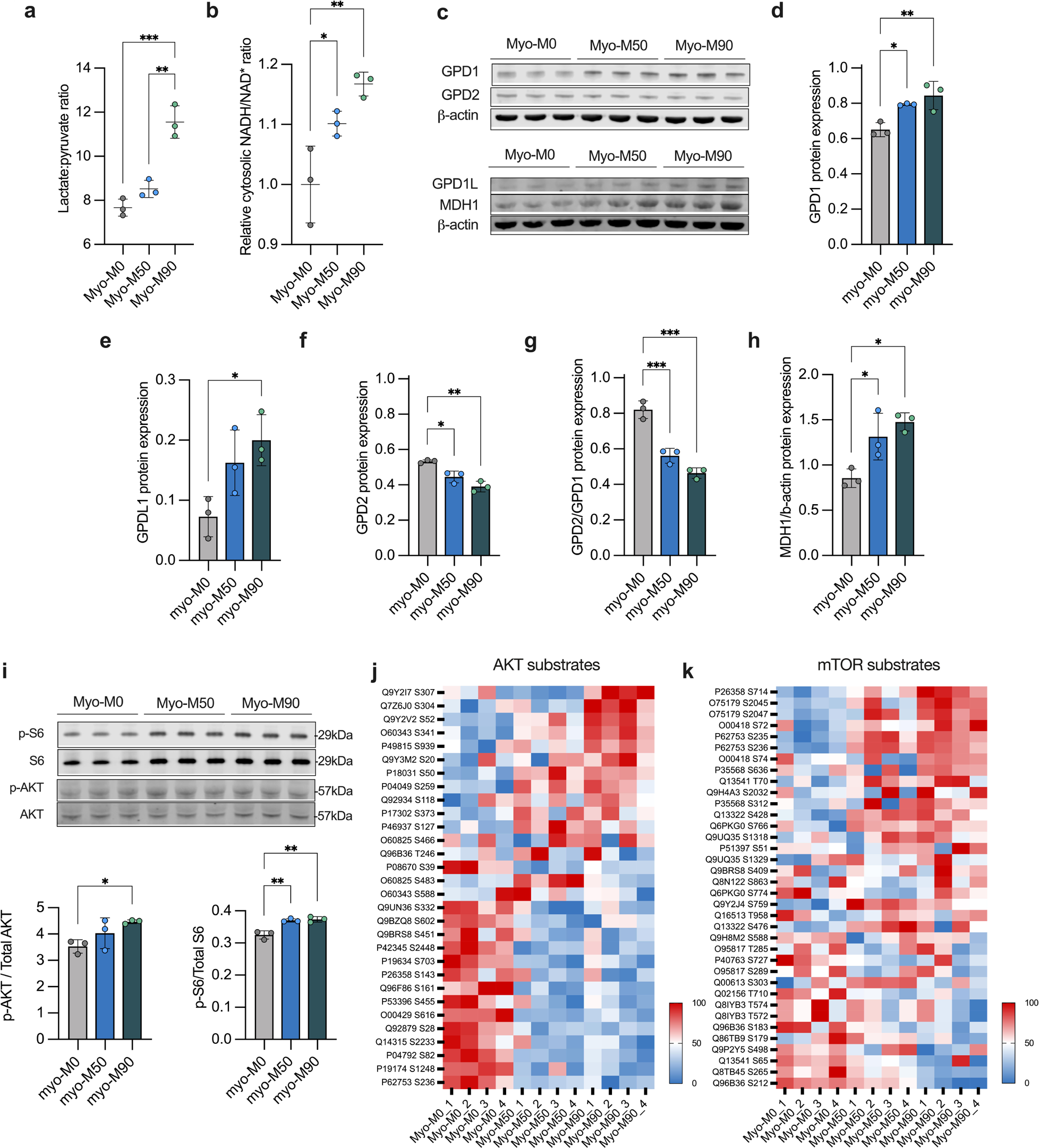
The m.3243A>G rewires cytosolic and mitochondrial metabolism. (A) Lactate to pyruvate ratio obtained from metabolomic analysis. n = 3 independent biological replicates. (B) Relative NADH/NAD ratio obtained from Peredox/mCherry measurements. n = 3 independent biological replicates. (C) Western blot of proteins associated to G3P and MA shuttles. (D, E, F and H) Quantification of proteins from (D). n = 3 independent biological replicates. (G) Ratio of protein expression of GPD2/GPD1. n = 3 independent biological replicates. (I) Representative western blot and quantification of phosphorylated S6 (S235/236) and AKT (S473) proteins. (J and K) AKT and mTOR substrates abundance obtained from phosphoproteomic dataset. n = 3 independent biological replicates. Source data are provided as a Source Data file. All data were represented as mean ± SD and data were analysed by one-way ANOVA with Tukey’s multiple comparisons test (*p < 0.05, **p < 0.01, ***p < 0.001, ****p < 0.0001).

As cytosolic NADH levels are also influenced by activity of the NADH shuttles, we measured the expression of proteins involved in the G3P (glycerol-3-phosphate) and malate-aspartate shuttles (Fig. 5c). These shuttles include enzymes such as cytosolic GPD1 (glycerol-3-phosphate dehydrogenase 1), GPD1L (Glycerol-3-phosphate dehydrogenase 1-like protein) and mitochondrial GPD2 (glycerol-3-phosphate dehydrogenase 2), which play a role in the oxidation of NADH produced during glycolysis. GPD1 utilises NADH generated by glyceraldehyde 3-phosphate dehydrogenase (GAPDH) to convert dihydroxyacetone phosphate (DHAP) into glycerol 3-phosphate. GPD2 then oxidises glycerol 3-phosphate back to DHAP while simultaneously reducing flavin adenine dinucleotide and facilitating electron flow through the ETC. The expression of cytosolic GPD1 and its isoform GPD1L increased in relation to the mutant load (Fig. 5d, e). Conversely, the mitochondrial component of the shuttle exhibited an inverted trend (Fig. 5f), decreasing with increased mutant load. Additionally, the ratio between GPD2 and GPD1, revealed a discrepancy in the expression of these enzymes between the mitochondrial and cytosolic compartments (Fig. 5g). We also quantified the expression of MDH1, the main component of the malate-aspartate shuttle. MDH1 was increased in the mutant lines, possibly as a compensatory mechanism in response to the increase NADH levels in the cytosol (Fig. 5h).

### Metabolic rewiring activates the PI3K/AKT/mTORC1 axis in myofibers carrying a high mutant load

We previously reported that in fibroblasts and muscle biopsies from patients carrying the m.3243A>G mutation, the PI3K/Akt/mTORC1 pathway was constitutively activated (Chung et al., 2021), apparently serving to sustain the m.3243A>G mutant load. We therefore assayed the phosphorylation of S6, (a downstream target of mTOR) and AKT (Fig. 5i). Consistent with our previous observations, we found that this signaling pathway was significantly upregulated especially in the cell line with the higher mutant load, as evidenced by increased phosphorylation of both AKT and S6 (Fig. 5i). In the Myo-M50 mutants, despite a slight increase in S6 phosphorylation, we did not observe any significant differences in AKT phosphorylation levels. To gain a deeper insight into pathway activation, we conducted a comparative phosphoproteomic analysis. AKT indirectly activates mTORC1 by inhibiting the tuberous sclerosis complex ½ (TSC1/2), a suppressor of mTORC1 activity. Our findings primarily identified increased phosphorylation of TSC2 (P49815) in Myo-M90, supporting the activation of AKT observed in the Myo-M90 on the western blot. Furthermore, the phosphoproteomic analysis indicated increased phosphorylation of S6 (P62753) at serine 235 and 236 (Figs. 5 j, k), indicating mTORC1 activation revealing that the m.3243A>G impacts in a different magnitude the nutrient-sensing pathway of this model.

## Discussion

In this work, we combined clonal expansion of hiPS cells, functional and omics assays to characterise the metabolic phenotype of hiPSC-derived myofibres carrying the m.3243A>G mutation. The m.3243A>G mutation is the most common mtDNA mutation and varies in frequency across different populations. This mutation can result in maternally inherited diabetes and deafness (MIDD) in some cases, as opposed to the syndrome MELAS. Notably, the m.3243A>G mutation exhibits variability in different cells and tissues of the same person in part due to the presence of mitochondrial heteroplasmy, where cells contain different copies of wild-type and mutated mtDNA (Durham, Samuels, Cree, & Chinnery, 2007) while maintaining a normal genomic DNA. In this study, we have demonstrated that three clonally expanded lines, despite sharing the same nuclear genetic background, exhibit variable mitochondrial deficiencies and compensations. This variability complicates our understanding of the disease phenotype.

The generation of stem cells carrying mtDNA mutations has been used to produce neurons (Hämäläinen et al., 2013; Klein Gunnewiek et al., 2020; Yokota, Hatakeyama, Ono, Kanazawa, & Goto, 2017), retinal pigment epithelial (Chichagova et al., 2017), cardiomyocytes (Ryytty et al., 2022; Yokota et al., 2017), neuronal organoids (Winanto, Khong, Soh, Fan, & Ng, 2020), endothelial cells (Pek et al., 2019), and other relevant cell types (Ryytty & Hämäläinen, 2023). Nevertheless, it has proven difficult to differentiate stem cells into muscle cells carrying mitochondrial mutations. To generate muscle contraction, this tissue demands a high ATP turnover, comprising a diverse range of cellular processes that activate depending on the intensity and duration of the contraction. Because the muscle stores of ATP are small, the muscle needs to derive energy from phosphocreatine and muscle glycogen breakdown, enabling substrate-level phosphorylation; and oxidative phosphorylation by utilizing reducing equivalents from carbohydrate and fat metabolism. In a system where mitochondrial ATP is compromised and high glycolytic rates are present, muscle dysfunction may occur, especially when the m.3243A>G mutation affects the majority of mitochondria.

We and others have successfully generated hiPSCs carrying mutations of mitochondrial DNA and the presence of the m.3243A>G mitochondrial DNA mutations does not interfere extensively with muscle cell reprogramming. This could be explained by the lower dependence observed in pluripotent cells on mitochondrial function. These cells switch to a more glycolytic phenotype, using glucose to sustain rapid cell proliferation and ATP production. Once reprogrammed, we showed that the mutant load remains stable over passage. Others have reported the existence of a potential bottleneck effect during the derivation of cells into the desirable cell type, resulting in a reduction of mutant copies of mtDNA and facilitating normal reprogramming (Chen & Guan, 2023). While our examination did not reveal significant changes in mutant load at these stages, a more detailed investigation into mitochondrial heteroplasmy during differentiation might be valuable. This approach would provide a comprehensive understanding of the dynamics involved in the differentiation process. Notably, despite the absence of major alterations in muscle differentiation, it is essential to highlight the presence of a proteomic signature indicating abnormal fibre distribution and morphology. It is acknowledged that the hiPSC-derived model does not represent a fully developed muscle model; instead, it constitutes a heterogeneous mixture of muscle fiber types. However, this model enables the measurement of key disease features specifically present in muscle tissue and identifies reproducible and robust readouts that could be used for drug screening and also identifies reproducible and robust readouts that could be used for drug screening. This underscores the significance of our findings in elucidating aspects of disease pathology that might be overlooked in more simplistic models.

While we have reported constitutive activation of the PI3K-AKT-mTOR pathway in fibroblasts, cybrid cells and muscle biopsies (Chung et al., 2021; Chung, Valdebenito, Chacko, & Duchen, 2022), it seems that this phenomenon might be cell type and mutant load specific, as we observed varying degrees of activation in the isogenic lines. Given that this pathway is a nutrient signaling pathway, our line with a mid-range mutant load appears to compensate for mitochondrial dysfunction by increasing tricarboxylic acid (TCA) activity and electron transport chain (ETC) proteins. It is important to note that the observations are not linear, as the Myo-M90 line exhibited different behaviour. Nevertheless, both cell lines were bioenergetically compromised, as evident in mitochondrial respiration and membrane potential results.

Variation in mtDNA heteroplasmy levels, specifically the 3243A>G mutation, has been described to lead to distinct phenotypic outcomes in patients, with mutant loads of 10-30% causing diabetes and occasional autism, 50-90% resulting in encephalomyopathies, and 90-100% leading to perinatal lethality. This has been attributed to transitions in cellular phenotype and gene expression, revealed through analyses of cybrids with increasing mutant mtDNA levels (Picard et al., 2014).

Overall, we have demonstrated the possibility of generating a muscle model derived from hiPSC expressing variable levels of the m.3243A>G mutation. This model showed that, regardless of the high levels of mutant DNA, the cells express a muscle phenotype characterized by contractility in response to chemical and electrical stimulation, given the formation of fully assembled sarcomeres. With this, we were able to explore mitochondrial dysfunction, where we observed changes in key mitochondrial variables, such as mitochondrial membrane potential and mitochondrial respiration. As expected, we obtained a higher mitochondrial NADH accumulation since the ETC is not able to fully oxidize this molecule, regulating TCA cycle flux. We also observed a major cytosolic NADH content in the mutant lines, suggesting an increased activity of NADH shuttles to replenish NAD+ content. Finally, these parameters affect nutrient sensing signaling pathways to a different degree, as reported in other cell models, which is corroborated with phosphoproteomics assays.

### Limitations of the study

In this study, two patient-derived cell lines and an isogenic control were studied in vitro. Due to the difficulties encountered during the reprogramming process of cells carrying mitochondrial DNA (mtDNA) mutations and their unstable nature in culture, it becomes challenging to obtain a broader range of induced pluripotent stem cell (hiPSC) lines. It is also worth noting that the model presented recapitulates development and generates myofibers. However, as of now, it has not been possible to generate fully mature muscle fibres, which also makes it impossible to compare our results with other cell models of the m.3243A>G mutation. Furthermore, no mouse model of this specific mutation has been created. Moreover, considering the variation in mutant load and phenotype from cell to cell as shown by others, a single-cell analysis would be valuable.

Long-term treatments are also challenging in a muscle model since the cells remain stable in culture for only one or two weeks. After this period, the cell matrix is unable to maintain the cell population, as spontaneous contractions occur, leading to cell detachment from the dishes.

## Acknowledgments

We thank Lu Yan and Olivier Pourquié for the muscle differentiation protocol. We thank Riccardo Zenezini and the UCL Mass-Spectrometry Science Technology Platform for the proteomic and phosphoproteomic analysis. We acknowledge the metabolic flux analysis facility of the Barts School of Medicine and Dentistry created with the support of the Barts and the London Charity - grant number MGU0401. We acknowledge Dr Monika Madej for the generation of the hiPSC clones. F.S.T. acknowledge support of the European Research Council (759108 – HISTOID). R.D.S.P. is funded by The Lily Foundation, Muscular Dystrophy UK (MDUK), and a seedcorn award from the Rosetrees Trust and Stoneygate Foundation. R.D.S.P. is supported by a Medical Research Council (UK) Clinician Scientist Fellowship (MR/S002065/1), Medical Research Council (UK) award MC_PC_21046 to establish a National Mouse Genetics Network Mitochondria Cluster (MitoCluster), and the LifeArc Centre to Treat Mitochondrial Diseases (LAC-TreatMito). R.D.S.P. and H.H. are supported by Medical Research Council (UK) strategic award MR/S005021/1 to establish an International Centre for Genomic Medicine in Neuromuscular Diseases (ICGNMD). The University College London Hospitals/University College London Queen Square Institute of Neurology sequencing facility receives a proportion of funding from the Department of Health’s National Institute for Health Research Biomedical Research Centres funding scheme. The clinical and diagnostic ‘Rare Mitochondrial Disorders’ Service in London is funded by the UK NHS Highly Specialised Commissioners. Early work in this area was supported by funding from Action Medical Research. A.R.C. is supported by the Medical Research Council, MRC DTP-iCASE programme [MR/RO15759/1]. G.E.V is supported by the National Agency for Research and Development (ANID)/Scholarship Programme/DOCTORADO BECAS CHILE/2019 – 7220052.

## Author contribution

G.E.V. and M.R.D conceived the project, designed and performed experiments. A.R.C, C-Y.C., P.S and B.O. provided resources and performed experiments. H.R, V.M. and K.B. performed and analyzed metabolomic data. H.H., F.S.T. and R.D.S.P. provided expert input on experimental design and analysis. G.E.V. and M.R.D. wrote the manuscript. All authors reviewed and approved the final version.

---

**Supplementary Figure 1. Maintenance of pluripotency and mitochondrial function during the undifferentiated stage of hiPS cells.**

(A) Representative images of hiPS cell colonies stained for nuclear pluripotency markers SOX2, NANOG and OCT4. Scale bar = 100 µm.

(B) Representative images of hiPS cell colonies stained for the pluripotency surface marker SSEA4. Scale bar = 100 µm.

(C) Relative gene expression of pluripotency markers thought RT-qPCR (n = 2 independent biological samples).

(D) Cross sectional fluorescence quantification of hiPSC colonies loaded with 25 nM TMRM. Coloured background represents 50 µM from the border of the colony to an arbitrary centre.

(E) Bright-field images of differentiating cells depicting the formation of myocentres. Arrows indicate the myocentres. Scale bar = 100 µm.

Source data are provided as a Source Data file. All data were represented as mean ± SD and data were analysed by one-way ANOVA with Tukey’s multiple comparisons test (*p < 0.05, **p < 0.01, ***p < 0.001, ****p < 0.0001).

**Supplementary Figure 2. Bundle formation and electrical stimulation in myofibres derived from hiPSC.**

(A) Representative confocal images of myofibres stained with antibodies against alpha-actinin 2 and pseudo-coloured in relation to the fibre orientation. Scale bar = 500 µM.

Representative changes in Indo-1 AM fluorescence intensity after electrical stimulation (0.5 Hz, 20 v). Stimulation is indicated by the arrow.

**Supplementary Figure 3. Fiber distribution in myofibres expressing the m.3243A>G**

(A) Comparison of the protein abundance obtained from the proteomic analysis. n = 3 independent replicates.

Source data are provided as a Source Data file. All data were represented as mean ± SD and data were analysed by one-way ANOVA with Tukey’s multiple comparisons test (*p < 0.05, **p < 0.01, ***p < 0.001, ****p < 0.0001).

**Supplementary Figure 4. Fiber distribution in myofibres expressing the m.3243A>G.**

(A) ARMS-qPCR of unrelated control and mutant lines expressing the m.3243A>G mutation. *n* = 3 independent biological replicates.

(B) Cell respiratory capacity measured using the Seahorse XFe96 extracellular flux analyser in myofibres (n = 3, 6 culture wells per experiment).

(C) Protein expression of mitochondrial respiratory complexes subunits (n = 3 independent biological replicates). TOM20 was used as loading control. Images are representative of at least three independent experiments.

(D) Representative confocal images of NAD(P)H autofluorescence.

Source data are provided as a Source Data file. All data were represented as mean ± SD and data were analysed by one-way ANOVA with Tukey’s multiple comparisons test (*p < 0.05, **p < 0.01, ***p < 0.001, ****p < 0.0001).

**Supplementary Figure 5. Targeted metabolomic analysis showed changes in labelling pattern of metabolites**

(A) Fractional enrichment of 13C isotopologues measured by targeted metabolomic (n = 3 independent biological replicates).sw

## Methods

### Maintenance of hiPSCs

Human induced pluripotent stem cells (hiPSCs) were cultured in mTeSR plus (Cat# 100-0276, StemCell Technologies) on Matrigel-coated plates (Cat# 354230, Corning Life Sciences) until they reached 70% confluency. Subsequently, cells were split using ReleSR (Cat# 100-0483, StemCell Technologies) in a 1:6 ratio. To prevent cell death, a 10 µM ROCK inhibitor (Cat# 1254/10, Bio-techne) was applied for one day. Daily media changes were performed, and the cultures were cleaned with a pipette tip under a microscope (EVOS™ XL Core Imaging System) to eliminate spontaneously differentiated cells that arose randomly around the edge of the colonies.

### Skeletal myogenic differentiation of hiPSCs

The direct reprogramming process was conducted based on an adaptation of Chal et al. Initially, hiPSC cultures were maintained in mTeSR plus (Cat# 100-0276, StemCell Technologies) on Matrigel-coated surfaces (Cat# 354230, Corning Life Sciences) until reaching 100% confluence before initiating myogenic differentiation. At this confluency, hiPSC cells were replated into an isolated cell suspension with 10 µM ROCK inhibitor (Cat# 1254/10, Bio-techne) at a very low density (no more than 12 cells per colony) and incubated overnight. The following day, the media was changed to remove the ROCK inhibitor and was maintained for 6 hours in in mTeSR plus (Cat# 100-0276, StemCell Technologies). The media was then replaced with DMEM/F12 (Cat# 11320033, ThermoFisher) supplemented with Insulin-Selenium-Transferrin (Cat# 41400045, ThermoFisher), 3 µM CHIR99021 (Cat# 4423/10, Bio-techne) and 500 nM LDN193189 (Cat# 04-0074-10, Generon). Media changes were performed every 2 days. On day 5, the media was supplemented with 20 ng/mL hFGF-2 (Cat# 450-33, PeproTech) for three additional days. Starting from day 6, the media was switched to DMEM/F12 supplemented with 15% KSR (Cat# 10828028, ThermoFisher), 10 ng/mL hepatocyte growth factor (Cat# 315-23, PeproTech), 2 ng/mL insulin-like growth factor-1 (Cat# 250-19, PeproTech), 20 ng/mL FGF-2 (Cat# 450-33, PeproTech), and 500 nM LDN193189 (Cat# 04-0074-10, Generon). From day 8, DMEM/F12 was supplemented with 15% KSR and 2 ng/mL IGF-1 until day 12. On day 12, the media was supplemented with 10 ng/ml HGF until day 22. Subsequently, cells were re-plated in Skeletal Muscle Growth Medium-2 (Cat# CC-3245, Lonza) for expansion. The medium was refreshed every two days until reaching 80% confluence. Myogenic progenitors were harvested and cryopreserved for downstream applications or replated for experimentation. All cultures were maintained in humidified air supplemented with 5% CO_2_ at 37 °C.

### Measurement of mtDNA mutant load

Levels of mtDNA mutation were detected using an allele refractory mutation system (ARMS)-based quantitative PCR (qPCR) analysis. DNA extractions from cells were performed using the DNeasy Blood & Tissue Kit (Cat# 69506, Qiagen). The concentrations of DNA samples were quantified using NanoDrop. Samples were diluted to 0.4 ng/μl. ARMS-qPCR primer working solutions (5 μM, 1 μL each; Forward [3243A]: CAGGGTTTGTTAAGATGGCAtA; Forward [3243G]: CAGGGTTTGTTAAGATGGCAtG; Reverse: TGGCCATGGGTATGTTGTTA) and SYBR Green JumpStart Taq Ready Mix (Cat# S4438, Sigma-Aldrich) were combined to create master mixes for mutant and wild-type genes. DNA samples (3 μL) and master mixes (7 μL) were pipetted into a 96-well PCR plate (Cat# MLL9651, Bio-Rad), and PCR amplification was performed using the CFX96 Touch Real-Time PCR Detection System (Bio-Rad). Each sample had three technical replicates. The mutant heteroplasmy level (%) was calculated using a previously described method, as shown in the equation below.

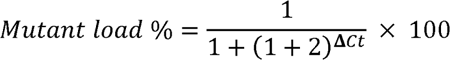

### RT–qPCR

RNA was extracted using the Qiagen RNAeasy Kit (Cat# 74104, Qiangen) following the manufacturer’s instructions. Reverse transcription was conducted in accordance with the Applied Biosystems guidelines. Gene expression analysis employed SYBR Green JumpStart Taq ReadyMix (Sigma-Aldrich #S4438) on the CFX-Connect RT–PCR System, utilizing CFX Manager Software version 2.1 (Bio-Rad) as per the manufacturer’s instructions. The qPCR data were analyzed using the delta-delta CT method. Primer sequences were obtained from Bono et al., 2021.

### Western blot

Myofibres were washed with ice-cold PBS once and lysed using 100 µl RIPA buffer (Cat# R0278, Sigma-Aldrich) with Protease and Phosphatase Inhibitor Cocktail (Cat# 78440, ThermoFisher). Cells were then scraped and stored at −80 C. Protein concentration was quantified using the Pierce BCA Assay Kit (Cat# 23227, ThermoFisher). For immunoblotting, 30 µg of protein samples in NuPAGE 4x LDS Sample Buffer (Cat# NP0007, Invitrogen) and 2% β-mercaptoethanol (Cat# 63689, Sigma-Aldrich) were boiled at 99°C for 5 min. Proteins were separated on 4-12% NuPAGE Bis-Tris polyacrylamide gels (Cat# NP0335, Invitrogen) and transferred onto PVDF membranes (Cat# IPFL00010, Millipore). Membranes were then incubated in Intercept (TBS) Blocking Buffer (Cat# 927-60001, Li-COR Biosciences) for 1 h at room temperature. After addition of primary antibodies diluted in the blocking buffer with 0.1% Tween-20, membranes were incubated overnight at 4°C on a shaker. Subsequently, membranes were incubated with appropriate secondary antibodies (Li-COR Biosciences; 1:10000; IRDye® 680RD Goat anti-Mouse IgG, #926-68070; IRDye® 800CW Goat anti-Rabbit IgG, #926-32211) for 1 h at room temperature before signals were developed with the LiCOR Odyssey CLx system.

### Mitochondrial membrane potential

Progenitors were seeded in 35 mm fluorodishes and differentiated until day 10. Myofibres were washed twice with the phenol-free recording medium (Cat# A1443001, Gibco) with 10 mM glucose, 1 mM glutamine, 10 mM HEPES, adjusted to pH 7.4; and then incubated with 25 nM tetramethylrhodamine methyl ester (TMRM) for 30 min at 37 °C. Cells were imaged with an LSM 880 (Carl Zeiss) confocal microscope using Fluar 63x/1.40 oil immersion objective lens at 37 °C. TMRM was excited with a 561 nm Argon laser with an output power of 0.2 mW. MBS 488/561 was used as a beam splitter and emitted fluorescence collected at 564-740 nm. Images were acquired using Zen Black software (Carl Zeiss) and fluorescence intensity was quantified using Fiji with the same threshold across all samples.

### The mitochondrial oxygen consumption rate

Measurements of oxygen consumption were conducted with the Seahorse Bioscience XFe96 bioanalyzer using the Seahorse XF Cell Mito Stress Test Kit (Cat# 103015-100, Agilent). hiPSC and progenitors were seeded on XF96 cell culture microplates (Cat# 102416-100, Agilent). On the day of the experiment, the culture medium was replaced with Seahorse XF Base medium (Cat# 103334-100, Agilent) supplemented with 1 mM pyruvate (Cat# 11360070, Gibco), 2 mM glutamine (Cat# 25030081, Gibco) and 10 mM glucose (Cat# A2494001, Gibco) and incubated for 30 min at 37 °C in a CO_2_-free incubator before loading into the Seahorse Analyser. After measuring basal respiration, the drugs oligomycin (1 µM), FCCP (1 µM, 2 µM), and rotenone/antimycin A (0.5 µM/0.5 µM) were added to each well in sequential order. Data were analysed using the XF Cell Mito Stress Test Report Generator. After the assay, protein was extracted from each well and a BCA assay was performed. The normalisation of the experiments is based on the relative protein obtained.

### Immunofluorescence

Pluripotent stem cells were seeded at single-cell colonies with ROCK inhibitor (Cat# 1254/10, Bio-techne) for one day. Then, cells were washed, and the media was changed daily until colonies appeared in the culture well. The cells were washed three times with 1X PBS (Cat# 14190144, ThermoFisher) and fixed in 4% paraformaldehyde for 15 min at room temperature and permeabilized with 0.1% Triton X-100 (Cat# 85111, ThermoFisher) for 30 min in PBS. The cells were then washed and incubated with primary antibodies (Myosin Skeletal Muscle antibody, MA1-90701; ACTN2 antibody, 14221-1-AP) in 3% BSA overnight at 4 degrees followed by incubation with Alexa Fluor-conjugated secondary antibodies for 1 h at room temperature. After antibody labelling, the coverslips were mounted on a glass slide using ProLong™ Gold Antifade Mounting (Cat# P36930, ThermoFisher) with DAPI and imaged using the confocal microscope as described above. Image post-processing was performed in ImageJ/FIJI.

### Mass spectrometry-based bulk proteomics and phosphoproteomic

Myofibres and progenitors were grown in 6-well plates. Culture wells were washed twice with PBS (Cat# 14190144, ThermoFisher) and subsequently RIPA buffer and 1X proteinase and phosphatase inhibitors was added directly to the plate. The cell lysate was collected by scraping the plate and boiled for an additional 10 min followed by micro tip probe sonication for 2 min with pulses of 1 s on and 1 s off at 80% amplitude. Protein concentration was estimated by BCA.

### Proteomic and phosphoprotemic data analysis

The original data was first log_2_ transformed and then only the proteins with at least 3 values from the 4 replicates were kept. At this point the missing data were imputed using automatic settings of Perseus. The proteomic dataset was then analysed using ExpressVis (Liu et al., 2022) and data visualization was done using GraphPrism and SRplot (Tang et al., 2023). The phosphoproteomic dataset was analyzed using Phosphomatics (Leeming et al., 2021) and data visualization was done using GraphPrism and SRplot (Tang et al., 2023).

### Calcium imaging

For imaging of cytoplasmic calcium, cells were washed once with phenol-free DMEM containing HEPES (Cat# 21063029, Gibco), then incubated with 1 μM Indo-1 AM (Cat# I1223, Invitrogen) and 0.02% Pluronic F-127 (Cat# P2443, Sigma-Aldrich) for 30 minutes at 37°C. Cells were then washed and incubated for a further 20 minutes in phenol-free DMEM at 37°Cs to allow de-esterification of intracellular AM esters. Cells were imaged using a UV-vis Zeiss LSM 880 confocal equipped with a 20× objective. Indo-1 fluorescence was excited at 355 nm, and emission measured simultaneously at 390 nm and 495 nm for Ca -bound Indo-1 and unbound Indo-1 respectively. For chemical stimulation, caffeine was dissolved in water and was added at a final concentration of 2.5 mM in recording media. For electrical stimulation, we used a MyoPacer Field Stimulator (IonOptix).

Images were analysed using ImageJ/FIJI. Regions of interest (ROI) were manually selected for each cell, and mean fluorescence intensity was quantified for all ROIs in each channel. Background was subtracted, and ratios between the emission signals of bound/unbound Indo-1 were calculated over time. The resulting ratioed traces representing cytosolic [Ca]c levels have been plotted.

### NAD(P)H autofluorescence

Progenitors were differentiated in 35 mm glass-bottom dishes. On the day of experiments, cells were washed twice with PBS (Cat# 14190144, ThermoFisher) and then incubated in recording media (DMEM no phenol red). Images were captured on a Zeiss 880 confocal microscope, equipped with a 60x UV-VIS oil immersion objective at 37 °C with excitation at 355 nm. Images were acquired under basal conditions, and after the addition of NaCN to a final concentration of 1 µM and FCCP at a final concentration of 1 µM. A wash with PBS (Cat# 14190144, ThermoFisher) was performed between the drug additions, and fresh media were replenished into the culture wells. The images were analyzed with Fiji, and the quantification was done following a previous publication (Chi, Bhosale, & Duchen, 2022).

### Lentivirus production

Peredox NADH/NAD+ sensor (Cat# 163060, Addgene) was transfected into HEK293 cells for lentivirus production. Briefly, the cells were seeded at 70-80% confluency in 10 cm dishes the day before transfection. The constructs (1.2 µg) were co-transfected with vesicular stomatitis virus G (600 ng) and pSPAX2 (800 ng) using Lipofectamine 3000 (Cat# L3000001, ThermoFisher). The lentiviral supernatants were collected 2 days after transfection, and then the cleared supernatant was concentrated with Lenti-X Concentrator (Cat# 631232, Takara) and resuspended in 1 mL of DMEM. Lentivirus transduction was evaluated using a CLARIOstar microplate reader (excitation/emission = 580/610 nm for mCherry, and excitation/emission = 400/513 nm for Peredox).

### Medium pH values

Medium pH values were measured based on the ratiometric property of phenol red. Progenitors were grown in 96-well plates with 150 µl of differentiation media and cultured for 5 days. On day 5, the media was replaced, and samples were collected on day 10. On the day of the experiments, the media from each well was transferred to a new 96-well plate, and the absorbance of phenol red at 443 and 570 nm was immediately measured. The higher the absorbance ratios of 443 to 570 nm, the more acidic the media. A BCA assay was done to ensure the same amount of cells/proteins were obtained per culture well.

### Targeted metabolomics

Progenitors were seeded in 60 mm dishes and differentiated for 10 days. Myofibres were then cultured in medium containing 5 mM C_6_-glucose (Cat# CLM-1396-5, Cambridge Isotope Laboratories) for an additional 18 h before metabolite isolation. Briefly, cells were washed with PBS (Cat# 14190144, ThermoFisher) three times and resuspended in ice-cold extraction buffer (20% ultrapure water, 50% methanol, 30% acetonitrile) at a ratio of 20 × 10^6 cells per ml. Subsequently, the cells were incubated on methanol and dry ice for 15 min, placed on a shaker for an additional 15 min at 4 °C, and then at –20 °C for 1 h. The cell lysate was centrifuged, and the supernatant was collected and transferred to autosampler glass vials, which were stored at –80 °C. LC-MS analysis was performed using a Q Exactive Quadrupole-Orbitrap mass spectrometer coupled to a Vanquish UHPLC system (Thermo Fisher Scientific). The liquid chromatography system was fitted with a Sequant ZIC-pHILIC column (150 mm × 2.1 mm) and guard column (20 mm × 2.1 mm) from Merck Millipore (Germany) and temperature maintained at 35°C. The sample (3 μL) was separated at a flow rate of 0.1 mL/min. The mobile phase was composed of 10 mM ammonium bicarbonate and 0.15% ammonium hydroxide in water (solvent A), and acetonitrile (solvent B). A linear gradient was applied by increasing the concentration of A from 20 to 80% within 22 min and then maintained for 7 minutes. The mass spectrometer was operated in full MS and polarity switching mode, in the range of 70-1000m/z and resolution 70000. Major ESI source settings were: spray voltage 3.5 kv, capillary temperature 275°C, sheath gas 35, auxiliary gas 5, AGC target 3e6, and maximum injection time 200 ms. For the targeted analysis, the acquired spectra were analyzed using XCalibur Qual Browser and XCalibur Quan Browser software (Thermo Scientific).

